# Multiomic characterization, early detection, and therapeutic targeting of myeloid sarcoma

**DOI:** 10.64898/2025.12.04.689069

**Authors:** Bettina Nadorp, Audrey Lasry, Sanam Loghavi, Ravi Patel, Hager Mansour, Benjamin J. Kelly, Christopher J. Walker, Jill Buss, Isaiah Boateng, Wafa Al-Santli, Zoe Ciantra, Rebecca Austin, Adrienne Heyrosa, Helee Desai, Andrea Laganson, Hasan Abaza, Linda Procell, Tejas Patel, Benjamin Kaffenberger, Saranga Wijeratne, Maria Guillamot, Maria Velegraki, Luis Chiriboga, Zihai Li, Lynne V Abruzzo, Daniel A. Pollyea, Christine McMahon, Arwa Shana’ah, John C. Byrd, Alan Shih, Ross L Levine, Eirini P. Papapetrou, Aristotelis Tsirigos, Elaine R. Mardis, Alice S. Mims, Iannis Aifantis, Ann-Kathrin Eisfeld

## Abstract

Myeloid sarcoma, an aggressive extramedullary subtype of acute myeloid leukemia (AML), occurs in ∼10% of patients, and has not yet been included in large-scale genomic studies. The critical biological changes that drive tumor evolution are unknown, its detection in asymptomatic patients remains a clinical challenge, and treatment options are limited as patients are often excluded from clinical trials, rendering it a neglected disease entity. Based on comprehensive multi-omic profiling, we demonstrate that myeloid sarcoma evolves from medullary AML with distinct sitespecific clonal evolution patterns. Additionally, we show that circulating tumor DNA sequencing can serve as a non-invasive method for molecular profiling of myeloid sarcoma, offering a novel avenue in molecular diagnostics. We characterize unique transcriptional profiles of myeloid sarcoma, reflecting immune evasion and adaptation to an extramedullary microenvironment. We provide evidence for a key role of RAS pathway activation and demonstrate in murine models of myeloid sarcoma that RAS inhibition effectively reduces tumor burden. Overall, our data highlight key differences between medullary AML and myeloid sarcoma including universal molecular evolution and RAS pathway activation as hallmarks of the disease and nominate RAS inhibition as a promising therapeutic strategy for patients with myeloid sarcoma.

## Introduction

Myeloid sarcoma is a distinct form of acute myeloid leukemia (AML) found in approximately 10% of patients with AML, in which myeloid blasts proliferate, expand, and form tumor-like lesions in extramedullary sites, which may present synchronously with AML or as isolated disease (1). Pathognomonic changes in the biology of the leukemic blasts that allow for tumor mass formation, including changes in the surrounding microenvironment, however, remain yet unknown. Myeloid sarcoma occurs with higher frequency in certain genetic subtypes of AML, including AML with monocytic features, favorable-risk subtypes such as NPM1-mutated and core binding factor AML (1). Unfortunately, patients with myeloid sarcoma have not benefited equally from recent advances in AML therapy, as they are routinely excluded from clinical trials. The true prognosis of patients with myeloid sarcoma is unclear given the inconsistent documentation of extramedullary disease and the suspected high numbers of undiagnosed myeloid sarcoma: for example, while testing for CNS involvement is recommended for high risk patients, imaging studies for possible myeloid sarcoma is guided by clinical symptoms. Moreover, the need for prompt imaging and tissue biopsy in newly diagnosed patients requiring treatment initiation can hinder early detection of myeloid sarcoma and subsequent treatment guidance. Thus, there is an unmet clinical need to detect extramedullary involvement and identify targetable vulnerabilities in myeloid sarcoma that, if functionally validated, may allow for the design of prospective clinical trials for this patient population.

We performed extensive transcriptomic sequencing at bulk, single-cell, and spatial levels, augmented by genomic profiling of extramedullary tumors and bone marrow (BM) samples from patients with myeloid sarcoma. Our data delineate the patterns of extramedullary clonal evolution of myeloid sarcoma, with a universally increased mutational burden, which is detectable in circulating tumor DNA (ctDNA) using plasma. We present evidence of RAS pathway addiction either through increased frequency of RAS pathway mutations or RAS/ERK pathway activation, independent of RAS pathway mutations, as a targetable vulnerability. We provide novel evidence of the efficacy of RAS inhibitors for the treatment of myeloid sarcoma in a murine model of disease. We provide novel insights into the pathogenesis of myeloid sarcoma, including the identification and characterization of gene programs associated with BM evasion and extramedullary homing with a gradual transition to a solid tumor environment.

## Results

### RAS mutations and pathway activation are cardinal features of myeloid sarcoma

To characterize the mutational landscape of myeloid sarcoma and associated medullary AML, we performed either paired tumor-normal whole-exome sequencing (WES, n=6) or targeted panel sequencing (n=36) on BM aspirates and myeloid sarcomas (**Figure 1A** and **Extended Data Table 1**). Myeloid sarcomas had a significantly higher tumor mutational burden (TMB) compared to matched medullary AML, with a median tumor mutation burden (TMB) of 1.66 vs. 0.64 mutations per megabase, respectively (p=0.0059, **Figure 1B**, **Extended Data Figure 1A**), as well as a higher burden of chromosomal copy number alterations (CNA; **Figure 1C, Extended Data Table 2**). Notably, alterations in the RAS pathway were the dominant molecular feature (43% of myeloid sarcoma, 48% of myeloid sarcoma BM, **Figure 1D**, **Extended Data Figure 1B** and **Extended Data Table 2**), which is substantially higher than the 15-25% reported general frequency of RAS pathway alterations in large-scale genomic studies of AML (3,4), or 28% in Alliance cohort patients without known extramedullary involvement (Fisher exact test, p=0.01252, **Figure 1E**). To confirm that mutations detected in myeloid sarcoma are of hematopoietic origin, we performed WES on myeloid sarcoma and paired flow-sorted BM blasts from three patients. In each patient we were able to detect myeloid sarcoma mutations in the sorted BM blasts (25.3%, 34.6%, and 50% in each patient respectively, myeloid sarcoma VAF>=5%, whole BM VAF<5%, see **Extended Data Figure 1C**, **Extended Data Table 2**), confirming that novel mutations detected in the myeloid sarcoma are of hematopoietic origin and are part of the myeloid sarcoma-forming clone.

**Figure 1.**
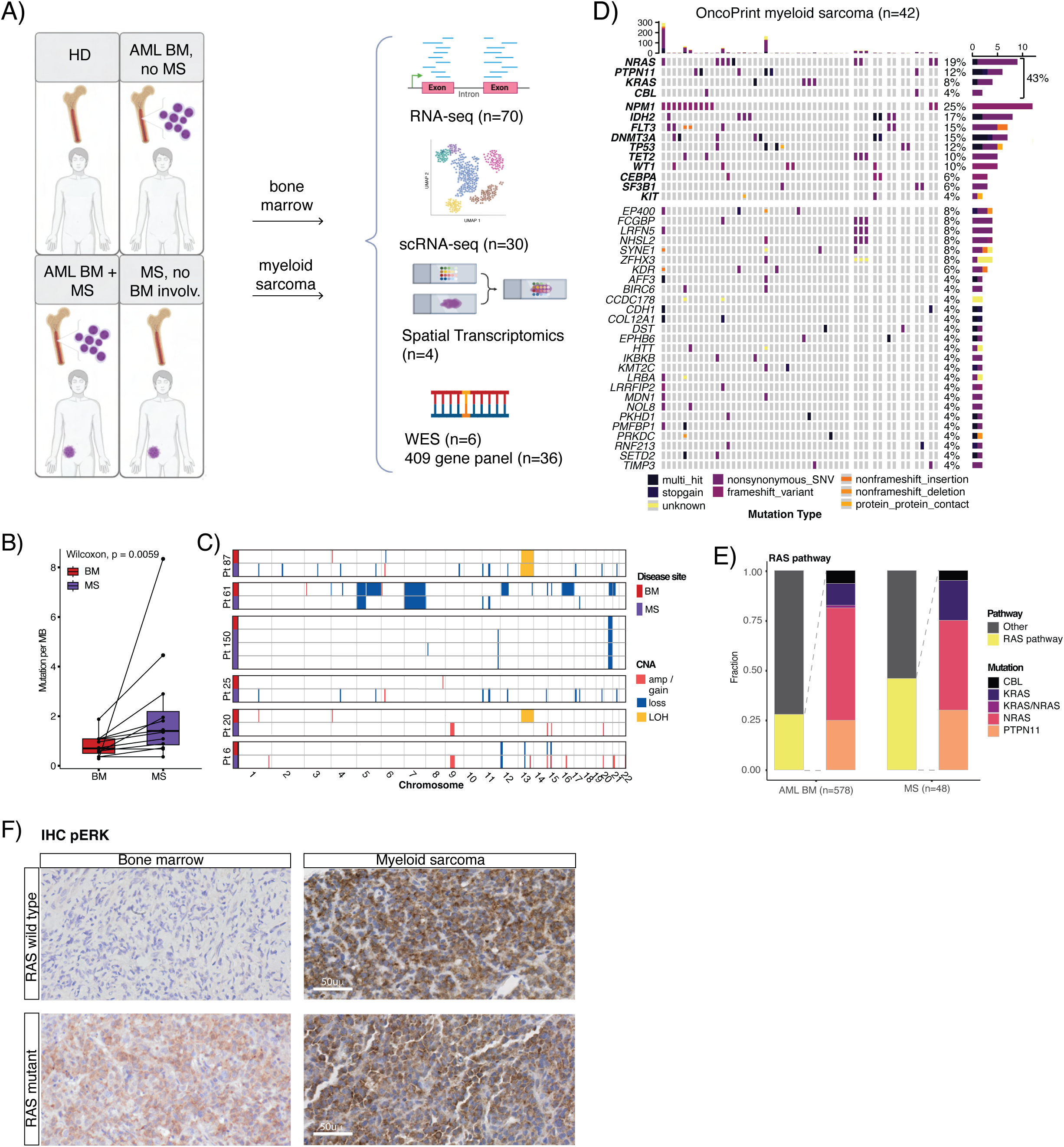
RAS pathway mutations are a cardinal feature of myeloid sarcoma. **A.** Experimental setup. **B.** Mutation burden in matched BM/myeloid sarcoma samples. A paired Wilcoxon test was used to evaluate statistical significance. Box plots represent the median with the box bounding the interquartile range (IQR) and whiskers showing the most extreme points within 1.5 × IQR. **C.** Chromosomal abnormalities detected in myeloid sarcoma in comparison to matched BM. MS: myeloid sarcoma, BM: bone marrow. **D.** OncoPrint of extramedullary disease mutations occurring in >1 patient (n=42 patients, n=48 myeloid sarcoma samples). Columns of patients with multiple myeloid sarcomas are separated to reflect patient count. **E.** Bar graphs reflect the fraction of RAS pathway mutations (CBL, KRAS, NRAS, PTPN11) in AML BM of patients without extramedullary involvement (n=578) compared to the mutational landscape in myeloid sarcoma (n=48). **F.** Acute myeloid leukemia in bone marrow showing p-ERK expression in a RAS pathway mutated case and lack of p-ERK expression in a RAS wild type case (IHC with hematoxylin counterstain, 100x); in contrast, extramedullary disease in 2 patients with RAS wild type and RAS mutant AML, respectively, both show expression of p-ERK, indicating RAS pathway signaling activation (IHC with hematoxylin counterstain, 100x).

Mutations in NRAS in our cohort included NRAS G13V (n=2), G12D (n=2), G12N (n=12), G12C (n=1) and G13D (n=1), while KRAS mutations included G13D (n=2), K117N (n=1), G6H (n=1), A146V (n=1) and V125A (n=1). Furthermore, KRAS mutations were more frequent in Alliance cohort patients with myeloid sarcoma compared to patients with AML (**Figure 1E**). The high prevalence of RAS pathway mutations suggested that activation of RAS signaling contributes to tissue evasion. To delineate RAS pathway activation in myeloid sarcoma patients with and without mutations in RAS pathway associated genes, we performed immunohistochemistry (IHC) for phospho-ERK (pERK) in BM and EMD of an additional cohort of 15 AML patients with myeloid sarcoma. In the BM, pERK was detected only in patients with RAS pathway mutations. Surprisingly, pERK was robustly expressed in extramedullary tumors of both RAS-mutant (8/9) and RAS-wild-type (5/5) myeloid sarcomas (**Figure 1F, Extended Data Table 3**). These data confirm that RAS pathway activation is a key component of myeloid sarcoma formation.

### Myeloid sarcoma is molecularly distinct and evolves from bone marrow AML

We then examined the clonal architecture of myeloid sarcoma and the associated medullary AML. We were able to detect linear and branched patterns of clonal evolution with unique myeloid sarcoma-specific mutations and CNA (**Figure 2A, Extended Data Figure 2A, and Extended Data Table 2**). In almost all cases, a shared parental clone was present in the BM, supporting the emergence of all extramedullary clones from a shared parental BM-based clone (**Figure 2A, Extended Data Figure 2A**). Moreover, we examined WES or targeted panel data from patients with multiple, synchronous, or metachronous myeloid sarcomas to compare their mutational profiles. We observed site-specific clonal evolution in synchronous (n=2 patients) and metachronous (n=5 patients) myeloid sarcomas, with a different dominant clone at each site (**Figure 2B, Extended Data Figure 2B**-**C**). Interestingly, in one patient with multisite involvement, mutations in all sites converged on the RAS pathway (**Figure 2B**). In contrast, in another patient, several mutations affected the MTOR pathway (**Extended Data Figure 2C**).

**Figure 2:**
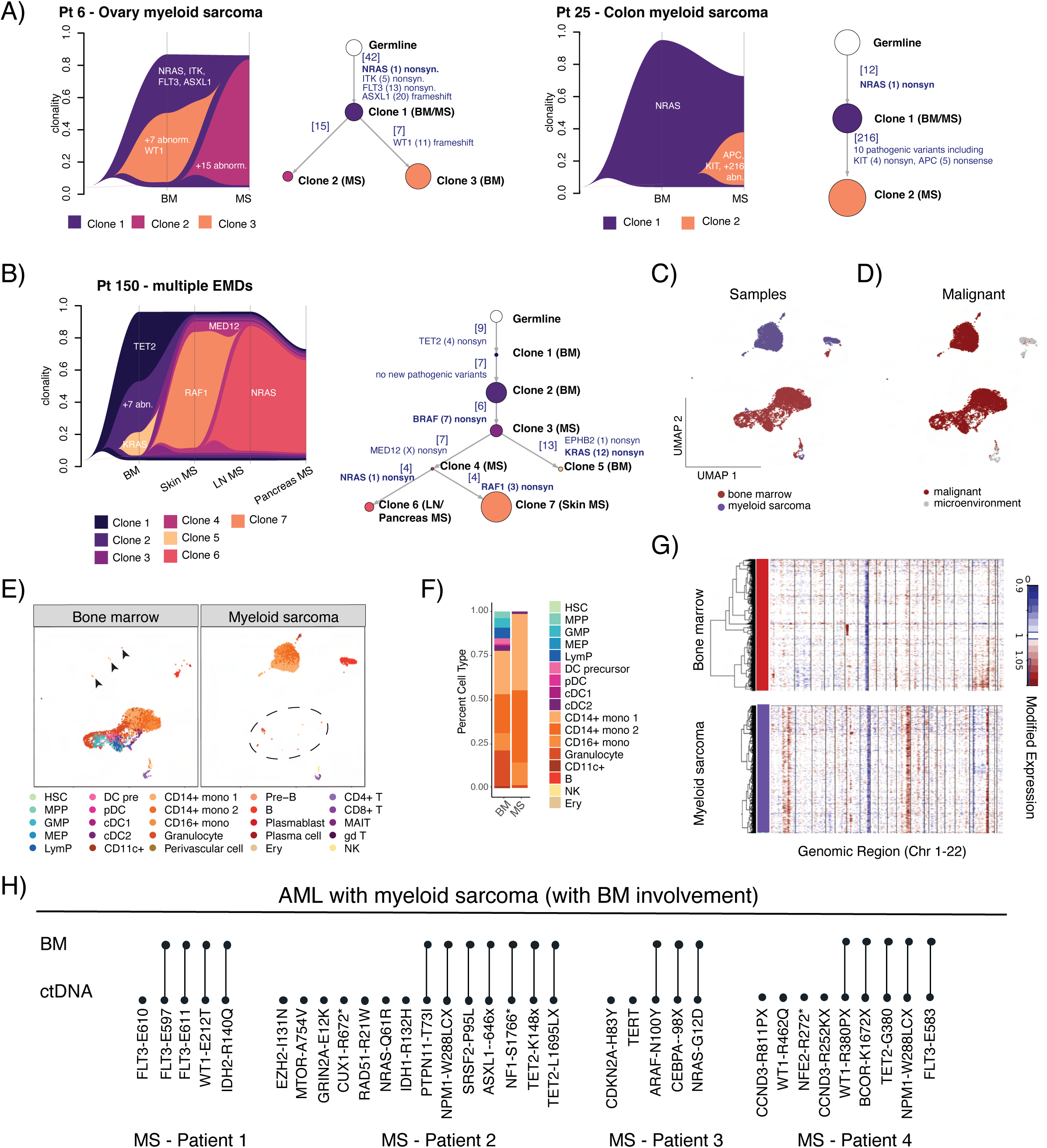
Myeloid sarcoma evolves from an ancestral BM clone through the acquisition of additional and unique genomic alterations. **A.** Clonal phylogeny and abundance of 2 matched myeloid sarcoma, BM pairs, comparing clones called in BM and myeloid sarcoma. [#] Mark the number of events (mutations or CNA) acquired in a clone. **B.** Clone abundance and phylogeny in a patient with synchronous myeloid sarcoma in different locations. **C.** Uniform manifold approximation and projection (UMAP) representing matched BM and myeloid sarcoma scRNA-Seq data of patient 3730, colored by location. **D.** UMAP colored by malignant cell annotation. **E.** UMAP split by location and colored by cell type reveals small proportions of myeloid sarcoma cells in BM (circled) and vice versa (arrowheads). **F.** Quantification of malignant cell subsets in BM vs. myeloid sarcoma. **G.** InferCNV heatmap of matched BM and myeloid sarcoma scRNA-seq data. MS: myeloid sarcoma, BM: bone marrow. **H.** Mutations detected in BM DNA or plasma ctDNA in patients with medullary AML and myeloid sarcoma demonstrating the presence of additional mutations in ctDNA, supporting the concept of clonal evolution of MS compared to BM-based AML..

To determine whether the clonal evolution of myeloid sarcoma results in transcriptomic and phenotypic changes, we performed scRNA-Seq on BM and myeloid sarcoma from a patient with concurrent medullary AML. We used inferCNV (5) to identify malignant cells and annotated cell populations using our previously published AML dataset (6) (**Figure 2C-2F**). Myeloid sarcoma malignant cells formed a distinct cluster from BM-based AML cells; however, unsupervised clustering revealed rare myeloid sarcoma-like cells within the BM and vice versa (**Figure 2E**). InferCNV analysis (5) further showed additional CNA in the myeloid sarcoma, validating the WES findings (**Figure 2G**). We next examined scRNA-Seq of matched BM and PB from 4 additional patients with concurrent medullary and extramedullary disease. Remarkably, in all cases, BM and PB cells clustered together, and we were unable to identify a myeloid sarcoma-specific population in the PB (**Extended Data Figure 3A-C**). These findings suggest that the clonal evolution of myeloid sarcoma occurs after it has homed to its final extramedullary destination rather than in the bone marrow or the blood circulation. This model is further supported by the site-specific evolution patterns observed in patients with synchronous or metachronous multi-site extramedullary involvement.

### Myeloid sarcoma can be detected using circulating tumor DNA

Early detection of myeloid sarcoma is challenging because, unlike patients with solid tumors or lymphoma, routine staging and surveillance with imaging are not performed in AML and no clear guidance for MS screening exists in current guidelines. Imaging studies and tissue biopsies are prompted only in patients with symptomatic manifestations indicative of extramedullary involvement. We hypothesized that myeloid sarcoma can be detected via plasma analysis of circulating tumor DNA (ctDNA), similar to the use of ctDNA for molecular profiling of other solid malignancies. Moreover, in light of its continued molecular evolution compared to BM-based AML, we speculated that the detection of a higher mutational burden in ctDNA compared to BM AML may indicate the presence of an additional extramedullary disease site. To test our hypothesis, we performed paired sequencing of myeloid sarcoma tissue and ctDNA obtained from plasma using a clinically validated and commercially available liquid biopsy assay. Mutations found in ctDNA corresponded to tissue biopsy (MS) results (**Extended Data Figure 3D**). These results serve as a proof of concept suggesting that ctDNA testing may be used as a surrogate for tumor tissue profiling. Furthermore, paired testing of BM-based AML and ctDNA from 4 patients with myeloid sarcoma showed the presence of additional mutations in ctDNA compared with BM (**Figure 2H**). In contrast, paired testing of BM and ctDNA of 3 AML patients without myeloid sarcoma showed identical mutations in ctDNA and BM samples (**Extended Data Figure 3E**). These results suggest that combining ctDNA with BM mutational testing may help identify patients with extramedullary disease and better profile the mutational landscape of the extramedullary disease component.

### Transcriptional and immune remodeling in myeloid sarcoma reflects adaptation to a solid tumor microenvironment

To examine the cellular composition of medullary and extramedullary AML at single cell resolution, we performed scRNA-Seq on myeloid sarcomas (n=3), BM from AML patients with and without myeloid sarcoma (AML+MS BM, n=8; AML BM, n=15, respectively), the BM of patients with myeloid sarcoma without overt BM involvement (MS BM, n=4) and healthy donor BM (HD, n=5). In two-dimensional space, each patient clustered separately, with some areas overlapping between different patients and healthy donors (**Extended Data Figure 4A-B**). We annotated malignant cells as described previously (6) (**Figure 3A, Extended Data Figure 4C-D,**) and compared the cellular composition of myeloid sarcoma and medullary AML in the different patients (**Extended Data Figure 4E**). Our analysis revealed that, in general, myeloid sarcoma had a cellular architecture reminiscent of medullary AML, consisting mainly of HSPC-like cells or more differentiated myeloid-like cells (**Extended Data Figure 4F**). However, the analyses of transcriptional profiles of malignant HSPC- and myeloid-like cells in our scRNA-Seq data (**Extended Data Figure 4G-H**) as well as on deconvoluted (8) malignant cells from our bulk RNASeq of AML and myeloid sarcoma (**Extended Data Figure 5A-B**) revealed significant differences between medullary AML and myeloid sarcoma, with myeloid sarcoma clustering separately from AML (**Figure 3C, Extended Data Figure 4G**). Comparison of differentially expressed genes in the scRNA-Seq data and the bulk RNA-Seq data allowed us to identify pathways that are uniquely upregulated in myeloid sarcoma compared to healthy BM, AML BM, or myeloid sarcoma-associated BM, including epithelial-mesenchymal transition (EMT), Interferon α (IFNα) and IFNγ response, and KRAS signaling (**Figure 3C-D, and Extended Data Table 4**). Surprisingly, while the HLA class II antigen processing machinery was downregulated in medullary AML, in myeloid sarcoma the levels of HLA class II antigen processing and presentation genes (*HLADMA/DMB/DRA/DOB, B2M*) were similar to those observed in healthy donor BM (**Figure 3E**), suggesting myeloid sarcoma employs different mechanisms of immune evasion compared to medullary disease. In addition, comparison of the transcriptional profiles of deconvoluted malignant cells from AML patients with and without myeloid sarcoma revealed that the malignant cells of AML patients with associated myeloid sarcoma already expressed higher levels of genes associated with EMT (*INHBA, COL7A1, COL1A1*), IFNα, and IFNγ response (*IFI44L, IFIT3, IFI27, ISG15, CCL2, STAT1*) in the BM. *MMP9*, a matrix metalloproteinase previously implicated in AML migration (9), was specifically upregulated in myeloid sarcoma-associated BM (**Figure 3F**, **Extended Data Figure 5C-D**). In contrast, comparison of myeloid sarcoma and associated BM revealed the downregulation of inflammatory programs, including interferon response genes (*CXCL8, IRF8/2, IFITM3),* and an up-regulation of EMT genes (*SNAI2*, *VCAN*, *EMP3, SNTB1*) in the malignant cells of the myeloid sarcoma (**Figure 3G, Extended Data Figure 5E-F** and **Extended Data Table 3**).

**Figure 3.**
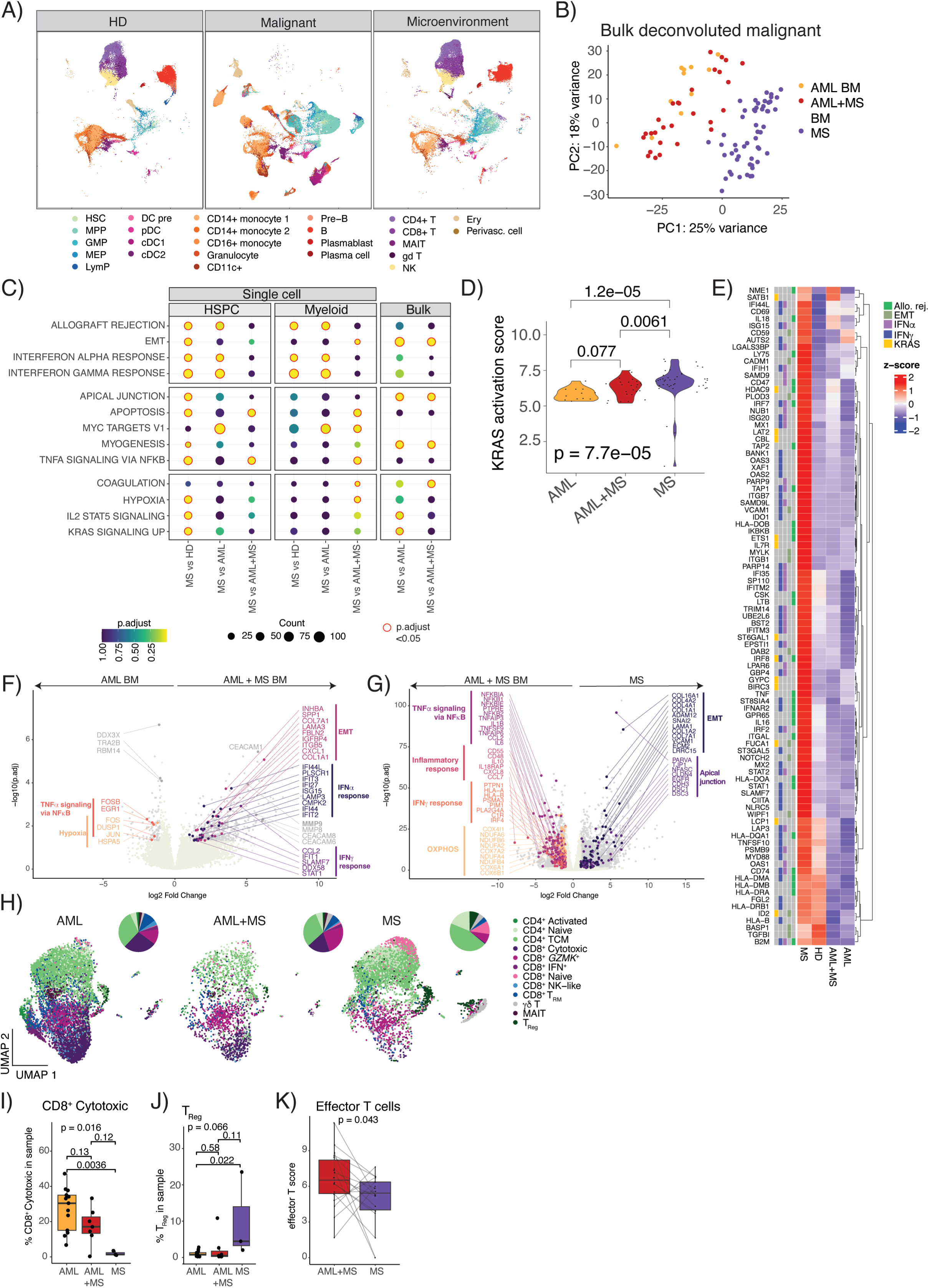
The transcriptional profile of myeloid sarcoma reflects transition from a “liquid” to a “solid” tumor state. **A**) Uniform manifold approximation and projection (UMAP) representing the healthy donors, malignant cells and microenvironment cells from AML BM,, AML+MS BM and myeloid sarcoma, colored by the cell type. **B.** Principal component analysis of BayesPrism deconvoluted malignant cell expression profiles of myeloid sarcoma (MS), associated BM (AML+MS BM), and AML BM without myeloid sarcoma (AML BM) from bulk RNA-seq data. **C.** Differential expression analysis comparing myeloid sarcoma (MS) to HD counterpart, AML BM with myeloid sarcoma (AML+MS) and AML BM without myeloid sarcoma (AML) hematopoietic stem and progenitor cells (HSPC) or myeloid cells in the single cell data, or MS to AML or AML+MS in bulk deconvoluted malignant cells. Dotplot summarizes the Hallmark pathway enrichment analysis in up-regulated genes in myeloid sarcoma. Only pathways enriched in at least two comparisons are shown. EMT: epithelial to mesenchymal transition. **D.** Violin plot showing enrichment of “KRAS activity up” Hallmark pathway in myeloid sarcoma (MS) compared to AML and AML+MS BM. Wilcoxon and Kruskal-Wallis tests were used to evaluate statistical significance. **E.** Average single-cell data group expression of differentially expressed genes in more than four comparisons from the “allograft rejection” (allo. rej.), “epithelial-mesenchymal transition” (EMT), “interferon alpha response” (IFNα), “interferon gamma response” (IFNγ), “KRAS signaling up” (KRAS) Hallmark pathways. The bottom rows annotate the pathway(s) to which each gene belongs; grey indicates the gene does not belong to the pathway. **F.** Differentially expressed genes between AML BMs with (AML+MS BM) and without (AML BM) myeloid sarcoma in bulk RNA-seq data. **G.** Differentially expressed genes comparing deconvoluted myeloid sarcoma and AML BM malignant cells of patients with myeloid sarcoma (AML+MS BM). **H.** UMAP representation of T cells from AML BM, AML+MS BM and MS. **I.** Quantification of CD8^+^ cytotoxic T cells in AML BM, AML+MS BM and myeloid sarcoma (MS). **J.** Quantification of T_Reg_ in AML BM, AML+MS BM and myeloid sarcoma (MS). **K.** Effector T cell scores in paired bulk RNA-Seq data of AML+MS BM and MS.

The prominence of inflammatory gene expression programs observed in myeloid sarcoma led us to examine immune responses in the BM and in extramedullary sites, as we have previously shown that in AML, inflammation is associated with an immunosuppressive microenvironment (6). Overall, T and B cell levels in AML, AML+MS BM and myeloid sarcoma were similar (**Extended Data Figure 6A-B**). However, analysis of the T cell compartment in our scRNA-Seq data (**Figure 3H, Extended Data Figure 6C**) revealed changes in the BM of patients with myeloid sarcoma as well as in the extramedullary site, with loss of cytotoxic T cells and an increase in regulatory T cells (T_Reg_) in extramedullary tumors (**Figure 3I-J**). In line with this, we observed lower effector T cell scores in myeloid sarcoma compared to paired AML BM in our bulk RNA-Seq data (**Figure 3K**), as well as lower effector T cell scores in myeloid sarcoma compared to AML and AML+MS BM in our scRNA-Seq data (**Extended Data Figure 6E**). Overall, these data indicate a unique transcriptional program in AML associated with myeloid sarcoma, characterized by genes essential for adaptation and growth within a solid tumor microenvironment. Furthermore, high levels of inflammation in the BM, along with suppressed immune response in extramedullary sites, may facilitate evasion of myeloid sarcoma cells from the BM and homing to extramedullary sites.

### Emergence of myeloid sarcoma leads to remodeling of the host microenvironment

To further characterize myeloid sarcoma in its host tissue, we used the 10X Visium spatial transcriptomics platform. To minimize the effects of different host tissues, we focused on one tissue, the skin. We examined four skin myeloid sarcomas compared to previously published spatial transcriptomics data on healthy skin (10). Examination of tissue architecture revealed significant remodeling of myeloid sarcoma-infiltrated skin as well as robust detection of malignant cells, using *SPN* (CD43), a highly sensitive marker for myeloid sarcoma (**Figure 4A-C, and Extended Data Figure 7A-B**). To further characterize the changes in the skin due to myeloid sarcoma infiltration, we used published scRNA-Seq datasets of healthy skin and calculated enrichment of the cell types captured by scRNA-Seq in our spatial data. Areas infiltrated by malignant cells showed enrichment of fibroblasts and endothelial cells, indicating vascularization and extracellular remodeling. Furthermore, infiltrated areas were enriched in immune cells, indicating a localized immune response to myeloid sarcoma (**Extended Data Figure 7C-F**). Notably, *SPN+* myeloid sarcoma infiltrated areas demonstrated high levels of the EMT-regulating transcription factors ZEB2 and SNAI2, as well as high levels of KRAS activation (**Figure 4D-E and Extended Data Figure 7A**). We further validated ZEB2 expression in myeloid sarcoma by immunohistochemistry on additional myeloid sarcoma samples from different tissues (**Figure 4F**). These data demonstrate both tissue remodeling in response to myeloid sarcoma emergence and the adaptation of myeloid sarcoma to the solid tissue microenvironment, with a critical role for RAS/ERK signaling in pathogenesis.

**Figure 4.**
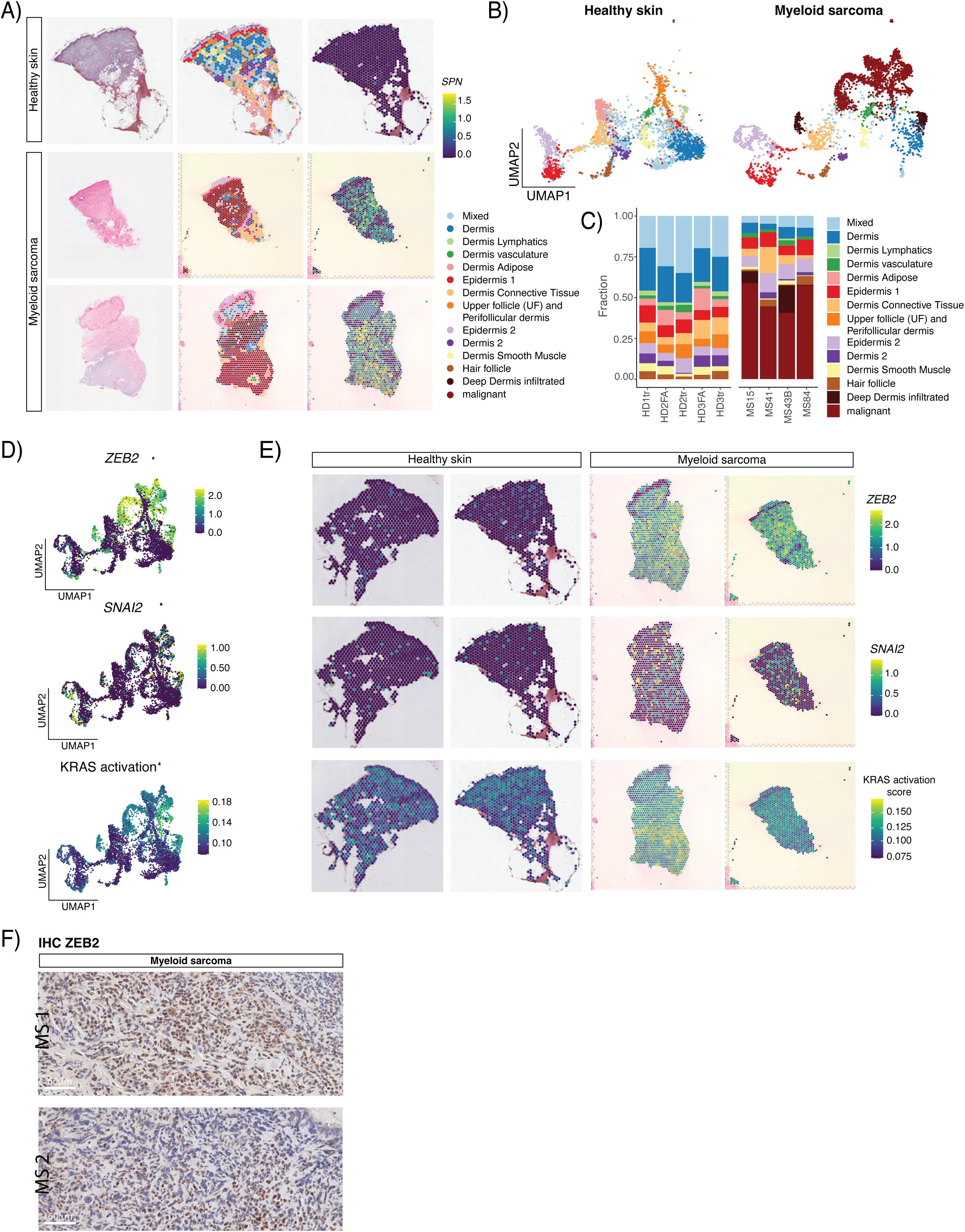
Myeloid sarcoma infiltration remodels the host microenvironment, activates RAS signaling and EMT-like programs. **A.** Representative Hematoxylin and Eosin stained image of healthy skin and skin of patients with myeloid sarcoma (left), representative spatial transcriptomics plot annotated by cluster/tissue region (middle), and SPN/CD43 expression marking the myeloid sarcoma (right). **B.** Split UMAP visualization of spatial spots annotated by cluster/spatial region. **C.** Quantification of cluster abundance in each patient. **D.** UMAPs show expression of epithelial-mesenchymal transition (EMT) transcription factors ZEB2 and SNAI2, and the KRAS activity Hallmark UCell score. **E.** Spatial transcriptomics plots showing ZEB2, SNAI2, and KRAS activity up Hallmark UCell score in healthy skin and skin with myeloid sarcoma. **F.** Myeloid sarcoma blasts showing expression of the EMT marker ZEB2 (Immunohistochemistry (IHC) with hematoxylin counterstain, 100x).

### RAS/ERK pathway inhibition reduces tumor burden murine models of myeloid sarcoma

To assess the role of RAS activation in myeloid sarcoma development *in vivo*, we took advantage of a genetically engineered human AML model with inducible NRAS^G12D^ expression, consisting of transplantation of CD34^+^ cord blood cells lentivirally transduced to express AML driver mutations, SRSF2^P95H^, ASXL1^del1900-1922^ and a doxycycline (Dox) inducible NRAS^G12D^ (SA+iR) into NSGS mice (12). These mice developed medullary and extramedullary AML involving the liver, spleen, kidney and lung (**Figure 5A-C**). To assess the role of RAS signaling we withdrew Dox treatment from one group of mice one week post-transplant (**Figure 5A**). In line with our mutational data, Dox-treated SA-iR mice had nodules on the liver and spleen, as well as myeloid infiltration into kidney and lung tissues, while SA-iR mice that were taken off Dox treatment did not show leukemic infiltration in these tissues (**Figure 5B-C, Extended Data Figure 8A**). Notably, infiltrating cells in Dox-treated mice expressed CD33, indicating myeloid differentiation (**Figure 5D**). These data show that human myeloid sarcoma is addicted to RAS pathway activation and suggest that therapeutic targeting of RAS signaling may be beneficial in the treatment of myeloid sarcoma patients.

**Figure 5.**
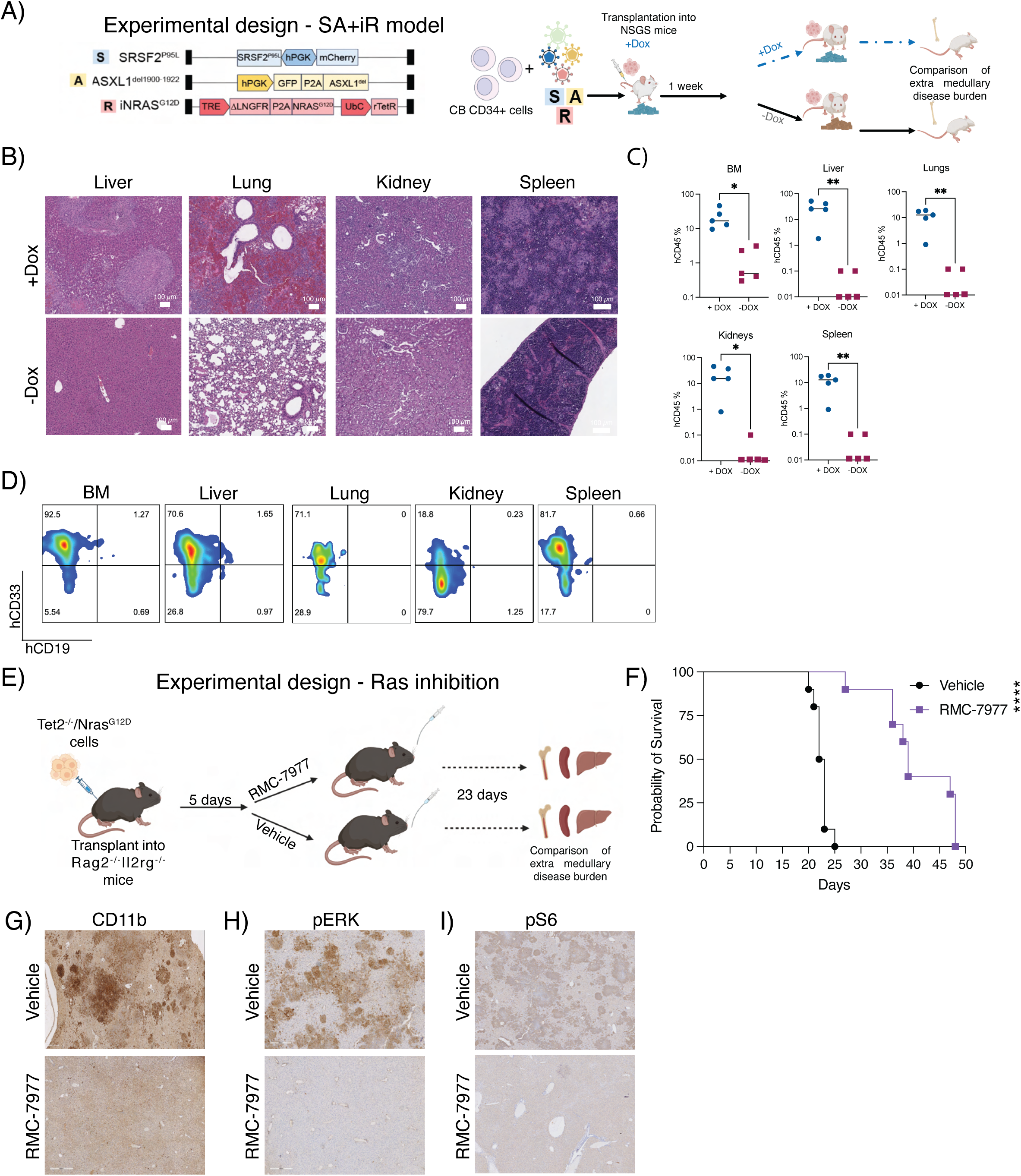
RAS pathway inhibition reduces tumor burden in a murine model of myeloid sarcoma. **A.** Schematic representation of SA-iR RAS on/off experiment. **B.** Hematoxylin and Eosin staining showing liver, lung, kidney and spleen in SA-iR +Dox (RAS on) and -Dox (RAS off) mice. **C.** Quantification of human CD45^+^ leukemic cells in different tissues in SA-iR +Dox and -Dox mice. Unpaired t-test, * p<0.05, ** p<0.005. **D.** Representative flow cytometry for CD33 and CD19 on human CD45 cells from SA-iR + Dox mice. **E.** Schematic representation of RAS inhibitor treatment in mice transplanted with TN cells. **F.** Overall survival of vehicle and RMC-7977 treated mice. Log-rank test, **** p<0.0001 **G.** CD11b IHC in livers of vehicle and RMC-7977 treated mice. **H.** pERK IHC in livers of vehicle andRMC-7977 treated mice. **I.** pS6 IHC in livers of vehicle and RMC7977 treated mice.

To confirm the role of RAS activation in myeloid sarcoma development, we established an additional murine model of myeloid sarcoma via transplantation of *Tet2*^-/-^/*Nras*^G12D^ (13) bone marrow cells into Rag2/Il2rg double knockout (DKO) recipients. We used immunodeficient mice, as in these hosts, Tet2^-/-^/Nras^G12D^ cells engraft without irradiation, which may alter the stromal niche in the BM and in extramedullary tissues and affect myeloid sarcoma development. Transplanted mice develop solid nodules on the spleen and liver, which were histopathologically confirmed to represent myeloid sarcoma (**Extended Data Figure 8B-C**). To examine the potential efficacy of RAS pathway inhibition in myeloid sarcoma, we transplanted Rag2/Il2rg DKO mice with *Tet2*^-/-^/*Nras*^G12D^ BM cells (13) and treated them with the RAS inhibitor RMC-7977 (**Figure 5E**), an oral and potent small molecule pan-RAS inhibitor, targeting both wild-type and mutated RAS proteins (14). RMC-7977-treated mice had increased overall survival compared to vehicle-treated mice (**Figure 5F**). To compare extra-medullary disease burden, we examined RMC-7977- and vehicle-treated mice three weeks post-transplant, when vehicle-treated mice started showing signs of disease. RMC-7977-treated mice had significantly smaller spleens compared to vehicle-treated mice (**Extended Data Figure 8D-E**)). Notably, while the spleens of vehicle-treated mice were heavily infiltrated by myeloid sarcoma, RMC-7977-treated mice had no or only focal residual myeloid sarcomas (**Extended Data Figure 8F-G**). Furthermore, the livers of vehicle-treated mice were extensively infiltrated by myeloid sarcoma, while the livers of RMC-7977-treated mice were free of tumor (**Extended Data Figure 8H**). Notably, myeloid sarcoma cells infiltrating the livers were positive for myeloid marker CD11b and expressed pERK and pS6 in vehicle-treated mice, indicating active RAS signaling in the tumors, while in RMC-7977 treated mice, the livers did not have aggregates of CD11b+, pERK+ or pS6+ cells (**Figure 5G-I**). Overall, our data indicates that RAS pathway inhibition is a promising approach for treatment of myeloid sarcoma.

## Discussion

Myeloid sarcoma is a distinct and aggressive myeloid neoplasm with a poorly understood molecular and biological basis and limited treatment options. Here, we present a cellular atlas of myeloid sarcoma and its associated medullary AML. We demonstrate that myeloid sarcoma has a high mutational burden and accumulates structural variants compared to associated BM-based AML, with inter-site heterogeneity of subclones in the same patient. This site-specific molecular evolution may explain the reported short-lived responses to targeted inhibitors (15), but may also represent an opportunity to screen for the presence of myeloid sarcoma via paired BM and ctDNA testing, in which a higher mutational burden with accumulation of subclonal mutations in ctDNA compared to BM may prompt imaging studies. While recognizing that these results must be validated in large, prospective patient cohorts, it is plausible to postulate that ctDNA testing may indeed be a reliable tool for the molecular profiling of myeloid sarcoma and may serve as a noninvasive surrogate for tissue biopsy in patients with myeloid sarcoma. This may be especially relevant if multiple myeloid sarcomas are present as it may serve as a sum of molecular aberrations present in medullary and extramedullary disease sites. Early detection of myeloid sarcoma is of high clinical relevance as it may enable early aggressive consolidation measures, such as allogeneic stem cell transplantation, as well as additional local treatments, as previously suggested in large-scale, retrospective studies (16).

We provide evidence for the cardinal role of RAS/ERK pathway activation in the pathogenesis of myeloid sarcoma, with robust enrichment of RAS pathway mutations compared with medullary AML (43% vs 28%) and mutation-independent pathway activation. We further demonstrate that mice transplanted with SA+iR (12) or *Tet2*^-/-^/*Nras*^G12D^ BM cells (13) develop myeloid sarcoma and that inhibition of the RAS pathway eradicates or reduces myeloid sarcoma burden in these mice. These findings nominate the RAS pathway as a key contributor to the pathogenesis of myeloid sarcoma, providing a rationale for investigating the potential efficacy of targeted RAS inhibitors in this clinically aggressive disease. Moreover, we characterize transcriptional programs in myeloid sarcoma, including activating pathways involved in EMT, which may be important for establishing a solid tumor mass in the extramedullary site, early signs of which can be detected in the BM of myeloid sarcoma patients. Interestingly, RAS pathway activation has been associated with EMT in solid tumors (17–19) suggesting the prevalence of alterations in this pathway in myeloid sarcoma may be directly associated with extramedullary colonization.

Together, our data suggest the use of ctDNA in the diagnostic workflow of myeloid sarcoma patients and establish the RAS pathway dependence of myeloid sarcoma. Our results provide preclinical evidence for therapeutic targeting of the RAS pathway, in combination with other AML-type therapies, as a treatment option for this aggressive disease that is often resistant to conventional cytotoxic chemotherapy.

## Methods

### Human patient samples

The OSU Leukemia Tissue Bank provided all cryopreserved bone marrow aspirates used in this study. All participants granted written consent for the storage and research utilization of their samples, adhering to the principles outlined in the Declaration of Helsinki and complying with the institutional review board regulation. The diagnosis of myeloid sarcoma and AML was confirmed by a retrospective review of hematoxylin and eosin-stained tissue sections from BM and extramedullary samples by an expert hematopathologist, according to WHO 2016 criteria (20)

#### Reference AML mutational profiles without extramedullary involvement

To characterize the mutational landscape of AML patients without myeloid sarcoma, we investigated the molecular characteristics of 872 adult patients with newly diagnosed AML enrolled on CALGB/Alliance study protocols based on intensive cytarabine/daunorubicin-based chemotherapy based on their reported myeloid sarcoma status. All patients gave written informed consent for participation in the studies. All study protocols were in accordance with the Declaration of Helsinki and approved by Institutional Review Boards at each treatment center. All patients were enrolled on CALGB 8461 (cytogenetic studies), CALGB 9665 (leukemia tissue bank), and CALGB 20202 (molecular studies) companion protocols. Mutational profiles of all patients have previously been published (4) and were performed centrally at the Ohio State University by targeted amplicon sequencing using the MiSeq platform (Illumina) and additional Sanger sequencing for CEBPA mutations, adding up to a total of 81 genes analyzed. **Figure 1C** only includes patients without extramedullary involvement.

#### Targeted mutation profiling

Targeted sequencing of myeloid sarcoma and extramedullary disease was performed using the Ion AmpliSeq Comprehensive Cancer Panel (Primbio), targeting the exons of 409 tumor suppressor genes and oncogenes encompassing over 50% of the Wellcome Trust Sanger Institute Cancer Gene Census (Suppl. Table 2).

#### Whole exome sequencing

##### Sorting of blasts

Cryopreserved BM samples were thawed in FACS buffer (PBS + 2% FBS) and stained with an antibody mix based on clinically annotated blast markers (DAPI, 1:3000, CD45-FITC 1:400, Biolegend, 368507, CD34-PE-Cy7, 1:400, Biolegend, 343515, CD11b-PE, 1:200, Biolegend, 301305). DAPI^neg^CD34^+^CD11b^+^ cells were collected in ice-cold FACS buffer and snap-frozen. For WES, DNA was extracted using the QIAGEN DNEasy blood and tissue kit, according to the manufacturer’s instructions.

##### Library preparation

Library construction for DNA extracts utilized the NEBNext Ultra II FS DNA Library Prep kit (New England Biolabs, Ipswich, MA) according to the manufacturer’s protocol. Whole exomes were captured using the xGen Exome Research Panel v2 and enhanced with the xGenCNV Backbone Panel and Cancer spike-in (Integrated DNA Technologies, Coralville, IA). Paired-end 151-bp reads were generated on the Illumina NovaSeq (Illumina, Inc., San Diego, CA).

##### Analysis

Secondary analysis was performed using Churchill (21), a comprehensive workflow for taking raw reads from alignment through to germline and somatic variant calls. Reads were aligned to the human genome reference sequence (build GRCh38) using BWA (v0.7.15). Sequence alignments were refined according to community-accepted guidelines for best practices (https://www.broadinstitute.org/gatk/guide/best-practices). Duplicate sequence reads were removed using samblaster-v.0.1.25, and base quality score recalibration was performed on the aligned sequence data using the Genome Analysis Toolkit (GATK) v4.1.9.0. Germline variants were called using GATK’s HaplotypeCaller. Somatic single nucleotide variation (SNV) and indel detection were performed using GATK’s MuTect-2 and VarScan. Somatic SNVs were filtered for quality (minimum site quality ≥100), population frequency (gnomAD maximum population frequency <0.0001), absence in normal data, somatic alternate read depth (inclusion if ≥4 reads), minimum tumor variant allele frequency (VAF) ≥5%, and genic location (within a coding or splice site (≤3-bp) region). Germline variation in cancer and other disease-associated genes and somatic variation across the coding region of the exome were analyzed. Copy number alteration (CNA), including loss of heterozygosity, was assessed using GATK’s CAN workflow and in-house-developed visualization. Clonal tracking across patient samples was performed using a customized version of the SuperFreq workflow (22) with adjusted input filtering and downstream processing and visualization scripts.

### Frozen human BM mononuclear cells preparation

Frozen human bone marrow samples were thawed and transferred into 50mL conical tubes containing PBS + 2% fetal bovine serum (FBS). Cell suspensions were centrifuged at 350 x g for 5 minutes at 4^°^C, and the supernatant was discarded. Samples were then subjected to dead cell depletion using a dead cell removal kit (Miltenyi Biotec, 130-090-101) or stained with DAPI (0.5mg/mL) and sorted for live cells (DAPI^low^), using a FACSAria IIu SORP cell sorter (BD Biosciences). All samples were gated based on forward and side scatter for cell sorting, followed by doublet exclusion, and then gated on DAPI^low^ for viable cells. Samples were sorted into 5mL poly-propylene tubes containing 300mL ice-cold PBS + 2% FBS. Following cell sorting, samples were centrifuged at 350 x g for 5 minutes at 4^°^C. For cell hashing, enriched live cells were tagged with cell-hashing oligo-tagged antibodies (Biolegend) according to the manufacturer’s instructions. Samples were then counted, and a maximum of 10^5^ cells for each sample were pooled together. For CITE-Seq, cells were pooled after hashing and stained with the TotalSeq-A human universal cocktail (Biolegend, 399907) according to the manufacturer’s instructions. Libraries were prepared using the Chromium Single Cell 3’ Reagent Kits (v3 and v3.1). Hashtag (HTO) libraries were prepared according to the NY Genome Center CITE-Seq and hashing protocol (https://citeseq.files.wordpress.com/2019/02/cite-seq_and_hashing_protocol_190213.pdf). Libraries were sequenced using an Illumina NovaSeq 6000.

### Single-cell RNA/CITE sequencing pre-processing

The Illumina bcl2fastq software was used to convert raw sequencing reads into FASTQ format. Subsequently, the Cell Ranger Single Cell Gene Expression Software (version 5.0, 10x Genomics) was employed to demultiplex and align the raw 3’ library reads to GRCh38 (version 2020-A). The Seurat R package (version 3.2.2) (23) was used for all downstream analysis, and visualizations were generated using ggplot2 (version 2_3.3.3). To eliminate low-quality cells and droplets that might have captured multiple cells, cells with less than 400 or more than 6000 unique feature counts and cells with over 15% transcripts originating from mitochondrial genes were excluded. This filtering process aimed to eliminate low-quality cells and droplets that might have captured multiple cells. Hashtag oligonucleotides (HTO) were normalized using a centered log ratio (CLR) transformation across cells, and HTODemux function in Seurat was applied to demultiplex hashed libraries (positive.quantile=0.99), and cells positive for more than one hashtag were excluded. We further used Souporcell (24) on hashed libraries to verify the patient identity and exclude doublets and cells in which genotype and hashtag assignment did not align. scDblFinder (version 1.5.13, https://github.com/plger/scDblFinder) was further used to identify doublets derived from the same patient in hashed and non-hashed libraries. Ambient RNA contamination was removed by applying SoupX (25)

The final dataset contained 169,777 cells, 19,084 cells from the BM of 5 healthy donors, 64,438 cells from the 15 AML patients without myeloid sarcoma, and 31,751 cells from the 8 AML patients with myeloid sarcoma, 4,403 cells from BM of patients with myeloid sarcoma, but not overt AML in the BM,12,606 cells from the peripheral blood of matched AML patients with myeloid sarcoma, and 30,884 cells from the MS. We captured 4054 cells per patient on average with a mean and median of 1669.395 and 1510 genes detected per cell.

RNA expression was normalized by total expression, multiplied by a scaling factor of 10,000, and then log-transformed. For Antibody Derived Tags (ADT), counts were center log-ratio transformed (divided by the geometric mean of each corresponding feature across cells and then logtransformed).

### Analysis of Single Cell RNA/CITE sequencing data

#### Clustering and visualization

To generate two-dimensional RNA expression visualizations, we performed principal component analysis (PCA) on the 2,000 most variable features and used uniform manifold approximation and projection (UMAP) (26) on the first 30 principal components (30 nearest neighbors, minimum distance 0.3). We then generated a shared nearest neighbor (SNN) graph with 30 nearest neighbors and subsequently clustered the graph (range of resolutions from 0.1-10). Resolution 2 (76 clusters) was chosen for downstream analysis.

#### Cell type annotation

To annotate cell types, we used the FindTransferAnchors function in Seurat (27) to identify the pairwise correspondence between cells in the annotated microenvironment/control cells manually annotated in (6) and all cells in this data set and projected the cell type labels using the TransferData function using the first 30 principal components (Broad_cell_identity). To annotate granular microenvironment cells, such as T, B, and NK cells, we subset to the individual compartments, integrated the data with Harmony (version 1.0) (28) and performed label transfer using the ‘Cell_type_identity’ annotations. Annotations were validated using differential expression analysis between cells labeled with a specific cell type against all other cells (Wilcoxon rank sum test with Bonferroni multiple-comparison correction, genes detected in at least 10% of the cell type cells, log2 fold change > 0.25 or < -0.25, adjusted p<0.05) and clusters with markers indicating contamination were removed.

#### Malignant and microenvironment division

Malignant and microenvironment cells were divided as in (6). In short, we used inferCNV (version 1.2.1) (29) to identify the malignant cells with clinically annotated abnormal karyotypes using the broad cellular compartments as annotations (HSPC, Myeloid, T, NK, B). We ran InferCNV with default settings except min_cell_per_gene=10, cutoff=0.1, denoise=TRUE, HMM=TRUE, analysis_mode=”subclusters”. We further validated the CNV+ T, B, and NK cells, running inferCNV on the T, B, and NK cells of all patients separately, annotating the more granular cell types within the broad cell type compartments and only kept CNV+ T, NK, and B cells that were detected in both analyses. Tumors without abnormal karyotypes were annotated using occupancy scoring (6) with a threshold of 0.7.

#### Differential gene expression

To identify differentially expressed genes between AML and AML with myeloid sarcoma (AML+MS), AML+MS and myeloid sarcoma, AML without myeloid sarcoma (AML) and myeloid sarcoma, or myeloid sarcoma and healthy donor cells, we broadly divided malignant cells or healthy counterparts into HSPC-like and Myeloid-like cells. To avoid strong patient-specific effects, we down-sampled the malignant cells to a maximum of 500 cells from each patient. To incorporate cellular detection rates, MAST (30), as implemented in Seurat, was used to perform differential expression analysis (genes detected in at least 10% cells, log2 fold change >0.25 or <-0.25, Bonferroni adjusted p < 0.05, and genes derived from X and Y chromosomes were removed due to male/female imbalance in the comparisons). Hallmark pathway enrichment was calculated using ClusterProfiler (31) and EnrichR (32) R packages. Gene set scores for Hallmark pathways ((33) were calculated using UCell ((34) as implemented in Seurat. To visualize commonly up-regulated genes across myeloid sarcoma comparisons, we plotted genes upregulated in 4 or more comparisons in the heatmap for Hallmark pathways occurring in four or more comparisons and three or more in Hallmark pathways occurring in fewer of the group comparisons.

## Immunohistochemistry

Immunohistochemistry (IHC) was performed on the Ventana Medical Systems Discovery Ultra platform using Ventana reagents. For Zeb2 IHC, we used Rabbit anti-human unconjugated Zinc Finger E-Box Binding Homeobox 2 (Zeb2), clone 1K22 (Protein Tech Group, catalog #82020, Lot#23004049: RRID: AB_2935598). After antigen retrieval in Cell Conditioner 1 for 32 minutes at 91°C, Zeb2 was diluted 1:50 (Ventana diluent Cat# 219) and incubated for 6 hours at room temperature, followed by goat anti-rabbit HRP Chromomap DAB detection. Human p-ERK IHC was done using the PERK (D11A8) rabbit mAb (Cell Signaling Technology) using a 1:300 dilution on a Leica-Bond iii automated stainer. Slides were counterstained with hematoxylin and mounted with synthetic media.

## Spatial transcriptomics

We used 10X Visium spatial transcriptomics platform (Visium CytAssist for FFPE Spatial Gene Expression 6.5mm, Human, 4 rxns [1000520], Dual Index Kit TS Set A, 96 rxn [1000251] and Visium CytAssist Spatial Gene Expression Reagent Kits, CG000495 RevE) to characterize the spatial gene expression patterns in four myeloid sarcomas in the skin. We followed all steps outlined in the 10X Genomics CytAssist Visium Protocol with overnight hybridization of ∼18h. Tissue preparation, staining, and High-Resolution Scanning were performed by OSUCCC CPDISR. Library generation, FFPE tissue total RNA quality, and sequencing of the resultant CytAssist Visium Spatial Transcriptomic libraries were performed by OSUCCC GSR. Healthy donor skin spatial transcriptomics samples were added from Castillo et al (35)

### Analysis

The Spaceranger Software (version 2.1.1, 10x Genomics) was employed to demultiplex and align the raw reads to GRCh38 (version 2020-A). The Seurat R package (version 4.3) (36) was used for all downstream analyses. Spots with low-depth coverage (<200 features/spot) were filtered. Individual samples were normalized using SCTransform, merged, and rpca integrated using default parameters with 3000 integration features and dimensions 1-30. 2D UMAP visualization was based on the 30 reciprocal PCA dimensions and 20 nearest neighbors. We then generated a shared nearest neighbor (SNN) graph with 20 nearest neighbors and subsequently clustered the graph (range of resolutions from 0.1-1.5). We used resolution 0.5 to annotate the spot using the healthy donor annotations from (37). We calculated the number of spots carrying labels from the healthy donors and spots derived from the myeloid sarcoma patients in each cluster and annotated the cluster based on the label with the maximum number. Clusters with the majority of spots derived from myeloid sarcoma patients were labeled malignant or infiltrated dermis based on hematopathologist input. We did not observe any spots from the upper follicle and perifollicular dermis in our myeloid sarcoma patients. Quantifications were visualized using ggplot2 (version 2.3.4.0).

We further performed multimodal integration analysis (38) to understand the cell type content in each spot using published single-cell RNA-seq data from the skin ((39,40) A hypergeometric test is used to compare the marker genes of each cluster to the marker genes of each cell type. Processed healthy skin single-cell data was downloaded from (41). We calculated cluster markers for the healthy donor skin and myeloid sarcoma skin spots separately and used the top 300 upregulated markers (adjusted p-value < 0.05, log fold change > 0.25) to calculate the enrichment score.

The Hallmark pathway (42) score for KRAS activity enrichment was calculated using UCell^41^ as implemented in Seurat.

### RNA-seq

#### Sample preparation

Tumor-derived RNA was subjected to 0.8x SPRI bead cleanup and size selection prior to DNase treatment and ribo-depletion prior to using the NEBNext Ultra II Directional kit preparation. The library was constructed for whole transcriptome sequencing (RNA-seq). Paired-end 151-bp reads were generated on the Illumina NovaSeq (Illumina, Inc.), and reads were aligned to the human genome reference sequence (GRCh38).

#### Data analysis

Transcripts per million (TPM) values were generated from paired-end RNA sequence data using Salmon with bootstrapping set to 100 (43). RNA-Seq-based diagnostic classification and outlier analysis was utilized following an analytical approach comparing samples to an internal Nationwide Children’s Hospital Institute for Genomic Medicine (IGM) pediatric cancer cohort and a cohort of pediatric and adolescent/young adult (AYA) tumors publicly available from the University of California Santa Cruz (UCSC) Treehouse Initiative (Compendium v11 and v9, https://treehousegenomics.soe.ucsc.edu/explore-our-data/). Salmon-derived TPM values for each sample were transformed using log2(TPM+1). Log2(TPM+1) values were obtained. Data were quantile normalized and filtered for known protein-coding genes within Biomart build GRCh38 (n=18,294). An n-of-1 differential expression analysis was performed using the following calculation: log2 fold change = log2 (gene A expression of the tumor sample) - log2(median gene A expression of the cohort). Mahalanobis distance and chi-squared *p-*value were used to determine statistical significance whereby outlier genes were classified as a log2 fold change >3.5 or <-3.5, and *P*<0.0005, similar to the methodology reported by Kothari et al (44)

### BayesPrism RNA-seq deconvolution

Raw RNA-seq reads were deconvoluted using BayesPrism (https://github.com/Danko-Lab/TED) (8) Due to expected tissue-specific cell types in myeloid sarcoma RNA-seq data, in addition to our own single-cell RNA-seq derived from the BM, PB, and MS, we added stromal cell types, as well as skin-specific cell types from the tabula sapiens human single-cell atlas (keratinocyte, pericyte cell, endothelial cell, epithelial cell, fibroblast, macrophage, vascular associated smooth muscle cell, stromal cell, and all cell types found in the skin) (45) Overlapping samples were used to validate deconvolutions while removing the patient single-cells from the reference data (**Extended Data Figure 5A**). Deconvoluted expression profiles of malignant cells were used in downstream differential expression analysis using DeSeq2 (46)

### Murine models

#### SAiR

Human cord blood CD34^+^ cells were transduced with lentiviral vectors encoding mutant SRSF2, ASXL1 and NRAS and transplanted into NSGS mice as previously described (12) One week post transplant, mice were randomized and one group was kept on Doxycycline chow (200mg/kg), while the other group was transferred to control chow. Mice were euthanized 6 weeks post transplant.

#### Tet2-/-/NrasG12D

Rag2/Il2rg double knockout (DKO) mice were obtained from Taconic Biosciences (model no. 4111) and housed at the New York University School of Medicine under pathogen-free conditions. All procedures were conducted in accordance with the *Guidelines for the Care and Use of Laboratory Animals* and were approved by the Institutional Animal Care and Use Committees at New York University School of Medicine. Viably frozen BM cells from Tet2^-/-^/Nras^G12D^ mice (13) were thawed and counted, and 100,000 cells/mouse were injected retro-orbitally into Rag2/Il2rg DKO mice. 5 days after transplantation, mice were randomized into RASi or vehicle treatment groups. RASi mice were treated with 10mg/kg of RMC-7977 (Selleckchem, E1858) as previously described (14).

### Data Availability Statement

All data generated in this study were submitted to the Gene Expression Omnibus (GEO) repository and can be accessed under GEO accession no. GSE241872. Additional AML BM and healthy control CITE-seq data were downloaded from GSE185381. Mouse WES data was downloaded from SRA project PRJNA833651.

## Acknowledgments

The authors are grateful to the patients who consented to participate in tissue banking and the families who supported them; to Christopher Manring and the CALGB/Alliance Leukemia Tissue Bank at The Ohio State University Comprehensive Cancer Center, Columbus, OH, for sample processing and storage services. The authors are especially grateful to R.C. and his family who donated his body to research and allowed for important scientific insights. Research reported in this publication was supported by the resources of the Pelotonia Institute for Immuno Oncology, an allocation of computing resources from The Ohio Supercomputer Center and Shared Resources (Leukemia Tissue Bank). The authors are grateful to Alan Shih and Ross Levine for sharing the *Tet2*^-/-^/*Nras*^G12D^ animal model. Research reported in this publication was supported in part by the National Cancer Institute of the National Institutes of Health under Award Numbers R01 CA262496, (A.K.E, E.R.M, A.S.M.), R01CA284595-01 (A.K.E., I.A.), R01CA283574-01 (A.K.E., E.R.M.), R01 LM013879 (A.K.E.), Leukemia & Lymphoma Society (A.K.E), the American Cancer Society (A.K.E.). We would like to thank the Genome Technology Center (GTC) for expert library preparation and sequencing and the Applied Bioinformatics Laboratories (ABL) for providing bioinformatics support and helping with the analysis and interpretation of the data. GTC and ABL are shared resources partially supported by the Cancer Center Support Grant P30CA016087 at the Laura and Isaac Perlmutter Cancer Center. The Aifantis Lab has been supported by the Vogelstein Family Foundation, the EvansMDS Foundation and the NIH/NCI (5 P01CA229086, 1 R01CA266212, R01CA271455, and 5 R01CA228135).

## Supplementary Figure Legends

**Supplementary Figure 1. Mutation Spectrum of extramedullary disease. A.** OncoPrint based on a 409-tumor mutation gene panel of six time-point matched myeloid sarcoma and associated AML BM samples. **B.** Oncoprint of AML bone marrow of patients with myeloid sarcoma. **C.** Variant allele frequencies (VAF) of myeloid sarcoma dominant mutations validated in sorted BM blasts.

**Supplementary Figure 2: Myeloid sarcoma clones exist at low VAF in the BM and show site-specific evolution. A.** Clonal abundance and phylogeny in matched BM and myeloid sarcoma samples (n=3). **B.** OncoPrint based on a 409-tumor mutation gene panel of myeloid sarcomas occurring in different locations or time points. **C.** Variant allele frequency of mutations detected in different myeloid sarcoma sites in a patient with synchronous multi-site myeloid sarcoma.

**Extended Data Figure 3: Circulating peripheral blood blasts resemble bone marrow disease in myeloid sarcoma patients. A.** UMAP representation of peripheral blood (PB, red) and associated BM (yellow) in 4 patients with myeloid sarcoma colored by sample and **B.** by cell type. Identified malignant cells are circled with dashed lines. **C.** Malignant cell type quantification in BM and PB. **D.** Mutations detected in DNA from myeloid sarcoma and paired plasma ctDNA. **E.** Mutations detected in BM DNA or plasma ctDNA in patients with AML without myeloid sarcoma.

**Extended Data Figure 4: Single cell landscape of AML and myeloid sarcoma. A.** UMAP of scRNA-Seq derived from BM of healthy donors (HD, n=5), AML (AML BM, n=15), AML with myeloid sarcoma (AMLMS BM, n=8) patients, and myeloid sarcoma patients without overt medullary disease (BM MS, n=4), matched peripheral blood of myeloid sarcoma patients (MS PB, n=5) and myeloid sarcomas (MS, n=3) Individual cells are colored by group annotation. **B.** UMAP as in A. colored by the donor. **C.** UMAP as in A. depicting CNV-positive cells as determined by inferCNV. **D.** UMAP, as in A., shows the occupancy score quantifying the fraction of patient cells within each cluster. **E.** Quantification of all cell types found in the BM and myeloid sarcoma. **F.** Quantification of malignant cell types found in medullary and extramedullary disease. HD: healthy donor, AML= AML BM, AML+MS: AML BM of patients with myeloid sarcoma, MS BM: BM of myeloid sarcoma patients without overt disease in the BM, MS: myeloid sarcoma, MS PB: BM matched PB of patients with myeloid sarcoma. **G.** UMAP representation of malignant healthy donor HSPC and myeloid cells and malignant cells from AML BM, AML+MS BM and myeloid sarcoma (MS). **H.** UMAP as in G, colored by cell type.

**Extended Data Figure 5: The transcriptional landscape of AML patients with myeloid sarcoma.**

A) Cell type quantification of whole BM of patient 26 as determined by scRNA-seq and BayesPrism bulk RNA-seq deconvolution, and malignant cells of Pt 119 myeloid sarcoma. B) Cell fractions determined by deconvolution of bulk RNA-Seq data with BayesPrism. C) Dotplot showing upregulated and D) downregulated pathways in BM of patients with myeloid sarcomas compared to AML without extramedullary involvement. HSPC and Myeloid reflect single-cell differential expression results, whereas bulk refers to the bulk RNA-sequencing data. E) Dotplot showing down-regulated Hallmark pathways across single-cell HSPC/Myeloid and bulk deconvoluted malignant cell differential expression analysis between myeloid sarcoma (MS) and its associated BM (AML+MS BM). F) Hallmark IFNα and IFNγ response scores in AML BM of patients with (red) and without myeloid sarcoma (yellow).

**Extended Data Figure 6: T cell response in myeloid sarcoma. A.** Quantification of T cells in AML BM, AML+MS BM and myeloid sarcoma (MS). **B.** Quantification of B cells in AML BM, AML+MS BM and MS. **C.** Quantification of T cell subsets in AML BM, AML+MS BM and MS. **D.** Effector T cell score in AML BM, AML+MS BM and MS.

**Extended Data Figure 7: Spatial remodeling of the skin in myeloid sarcoma.**

A) Spatial transcriptomics plots showing clustering, SPN/CD43, ZEB2, SNAI2 expression, and KRAS activation Hallmark UCell scoring on patients with myeloid sarcoma in the skin and healthy donor skin. B) Cluster quantification without malignant spots. C) Multivariable integration analysis on myeloid sarcoma and healthy skin clusters separately. Enrichment heatmaps of non-immune cell types from Reynolds et al.(47) and spatial clusters showing the adjusted p-value. Diff KC, differentiated keratinocytes; F, fibroblasts; LE, lymphatic endothelium; Prolif KC, proliferating keratinocytes; VE, vascular endothelium; KC, keratinocyte. D) Enrichment heatmap for immune cell types of Reynolds et al.(47). DC, dendritic cell; Macro, macrophage; moDC, monocyte-derived DC; MigDC, migratory DC; Mono, monocyte; Inf. mono, inflammatory monocyte; ILC, innate lymphoid cell; NK, natural killer cell; TC, cytotoxic CD8+ T cell; TH, CD4+ T helper cell. E) Enrichment of non-immune cell types and F) immune cell types from Hughes et al.(48)

**Extended Data Figure 8. RAS inhibition in a murine model of myeloid sarcoma**

A) Representative images of lungs, kidney, liver and spleen (from left ro right) from SA+iR +Dox and -Dox mice. B) Representative images of spleens (left) and liver (right) from Tet2^-/-^/Nras^G12D^ mice. C) Hematoxylin and Eosin staining (left) and CD11b IHC (right) on representative liver section from Tet2^-/-^/Nras^G12D^ transplanted mice. D) Representative pictures of spleens of vehicle or RMC-7977 treated mice. E). Spleen weights of vehicle and RMC-7977 treated mice. Unpaired t-test, **** p<0.0001 F). Representative H&E-stained sections of spleens from vehicle and RMC7977 treated mice showing extensive involvement by myeloid sarcoma in the vehicle-treated mice in comparison to only focal residual myeloid sarcoma (green circle) in the RMC-7977-treated mice. G) High magnification of myeloid sarcoma in vehicle-treated mice and uninvolved spleen and myeloid sarcoma from RMC-7977-treated mice. H) Representative H&E-stained sections of livers from vehicle and RMC-7977-treated mice showing extensive involvement by myeloid sarcoma in the vehicle-treated mice in comparison to no residual myeloid sarcoma in the RMC7977-treated mice.

