## Supplementary figures and images for "Multiomic characterization, early detection, and therapeutic targeting of myeloid sarcoma"

### Supplementary Figure 1

#### Extended Data Figure 1: Mutation Spectrum of myeloid sarcoma.

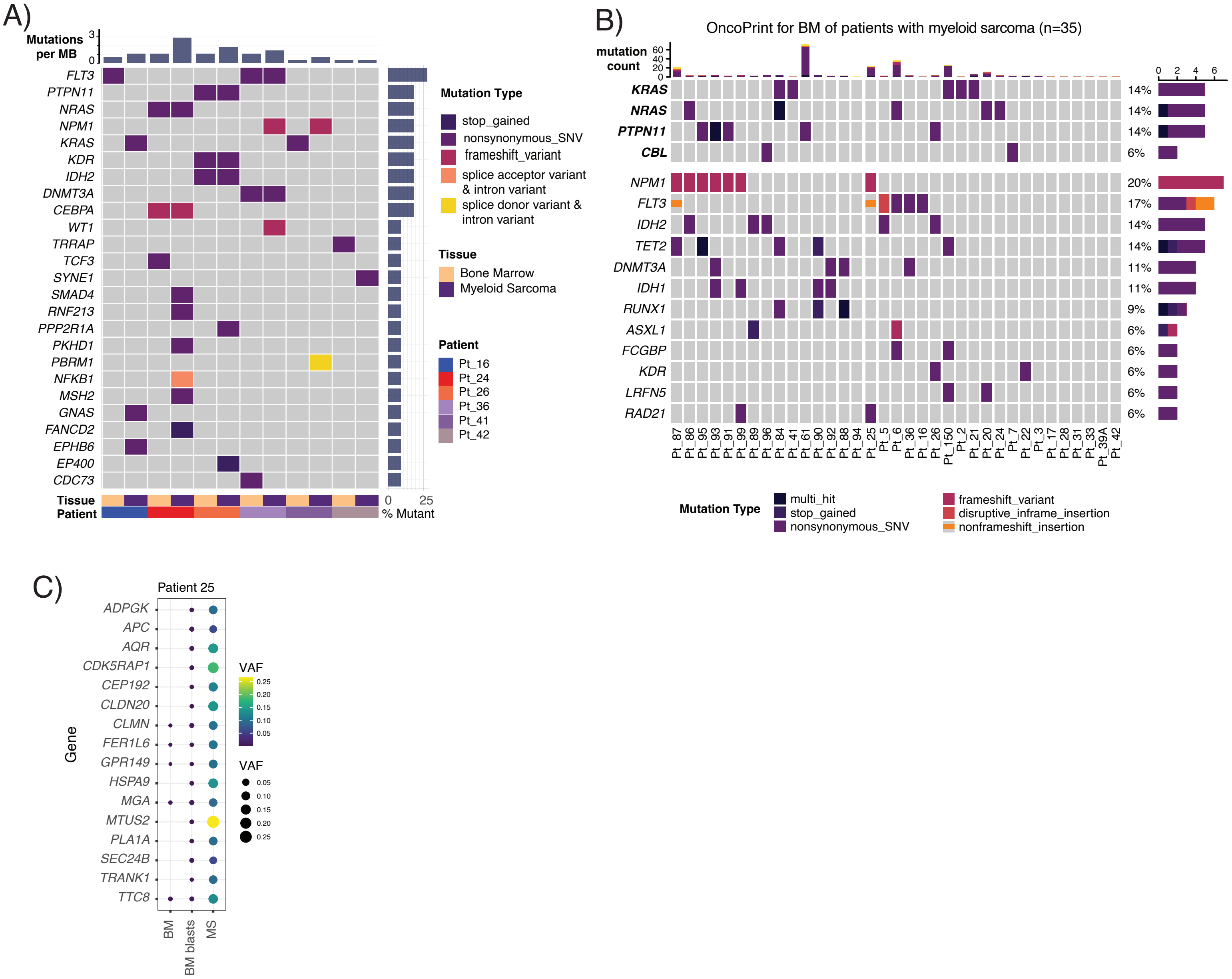

### Supplementary Figure 4

Extended Data Figure 4: Single cell landscape of AML and myeloid sarcoma.

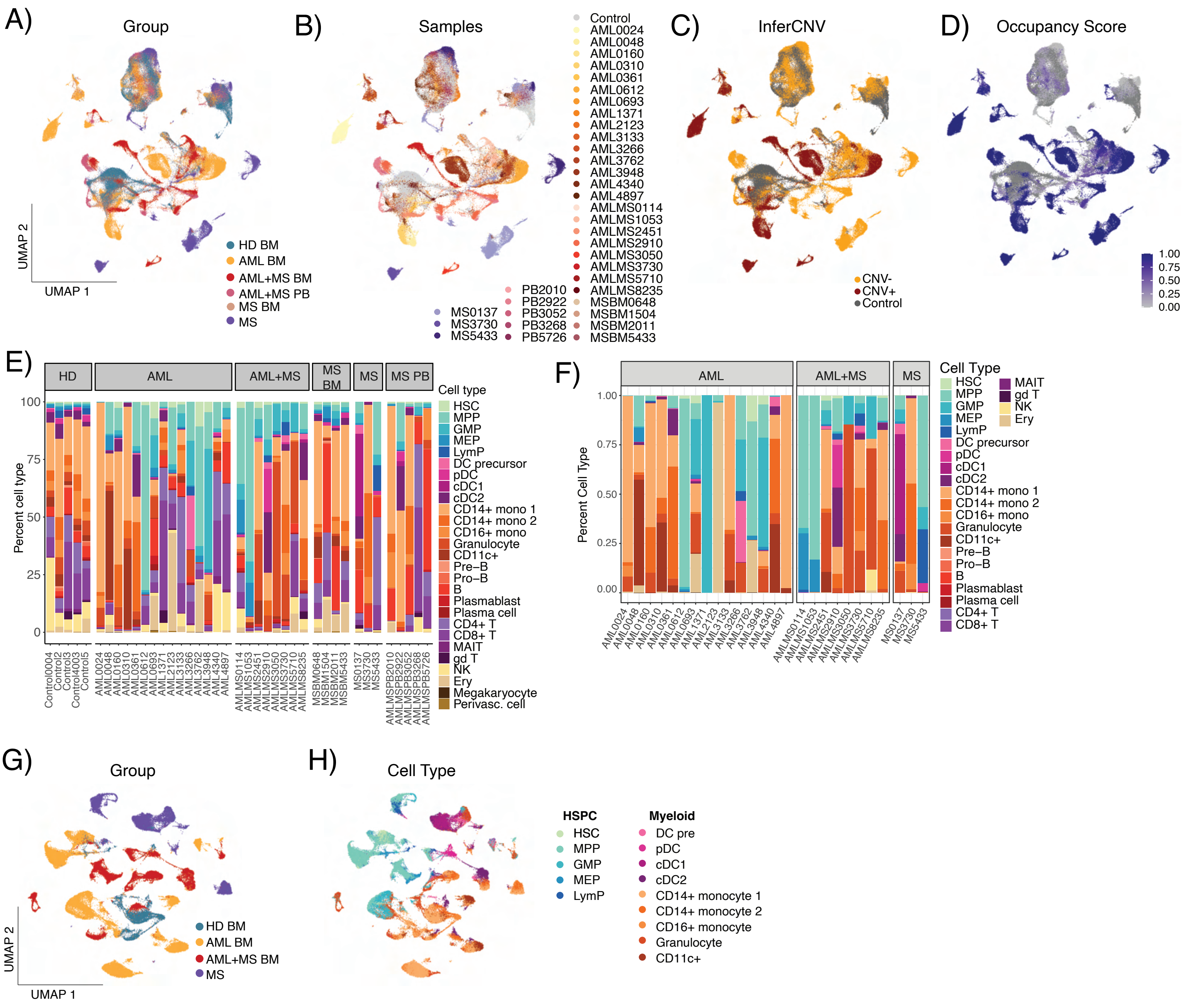

### Supplementary Figure 5

Extended Data Figure 5: The transcriptional landscape of AML patients with myeloid sarcoma.

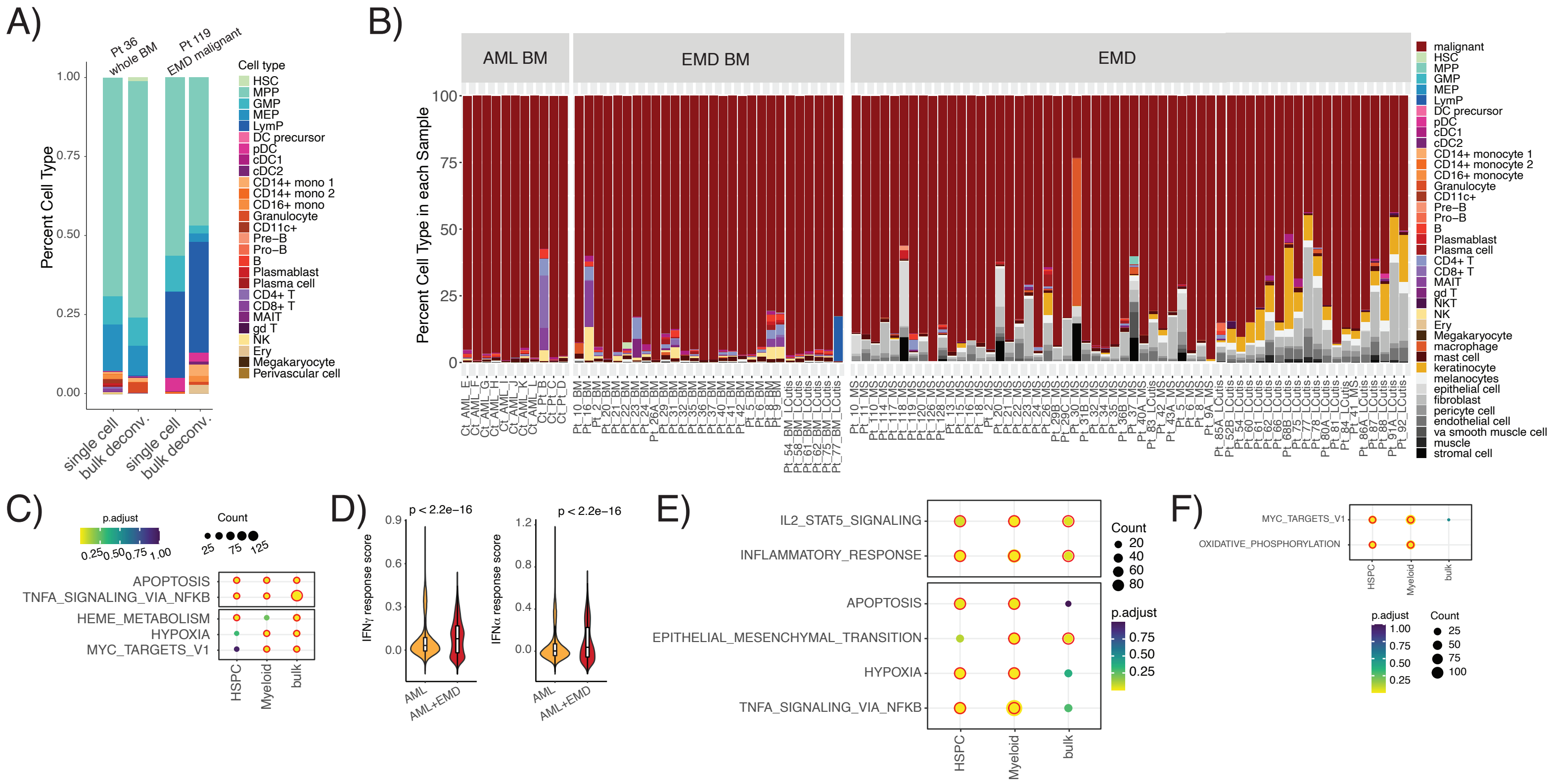

### Supplementary Figure 6

Extended Data Figure 6: T cell response in myeloid sarcoma

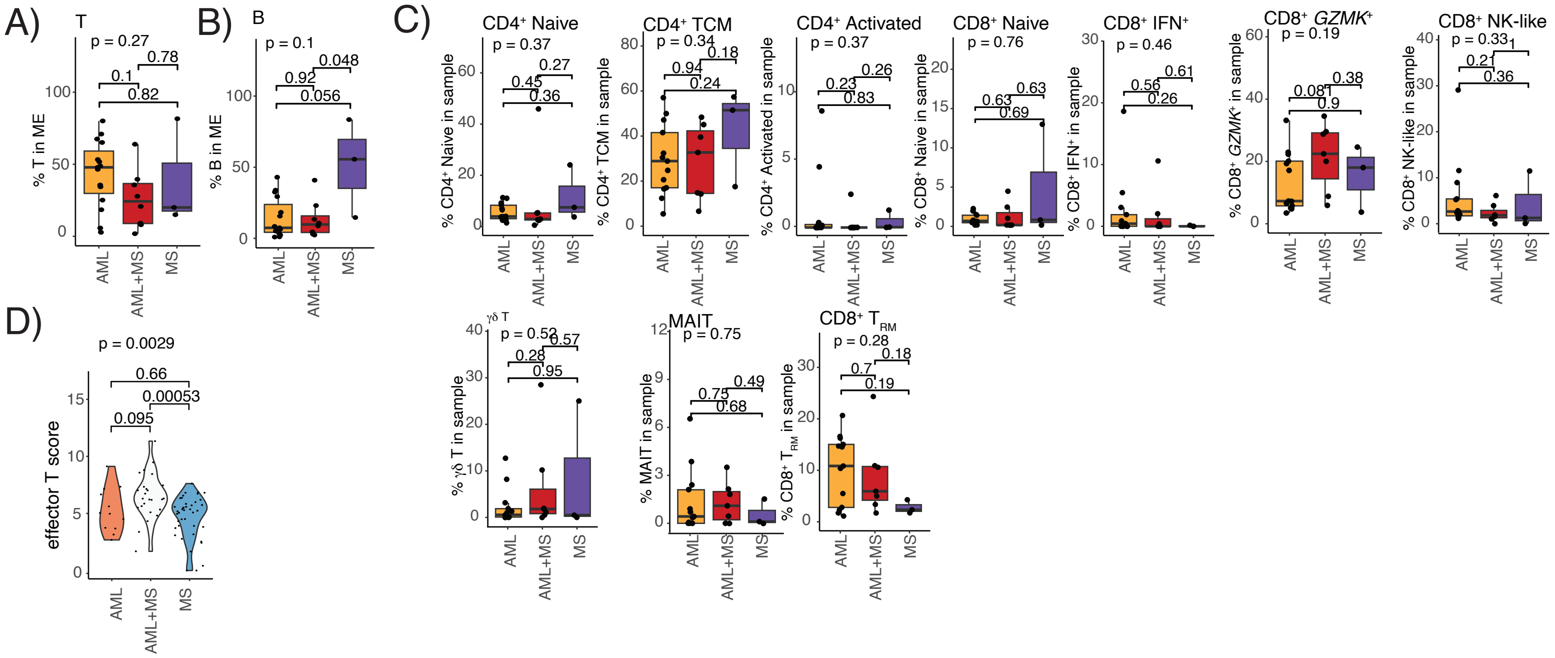

### Supplementary Figure 7

Extended Data Figure 7: Spatial remodeling of the skin in myeloid sarcoma.

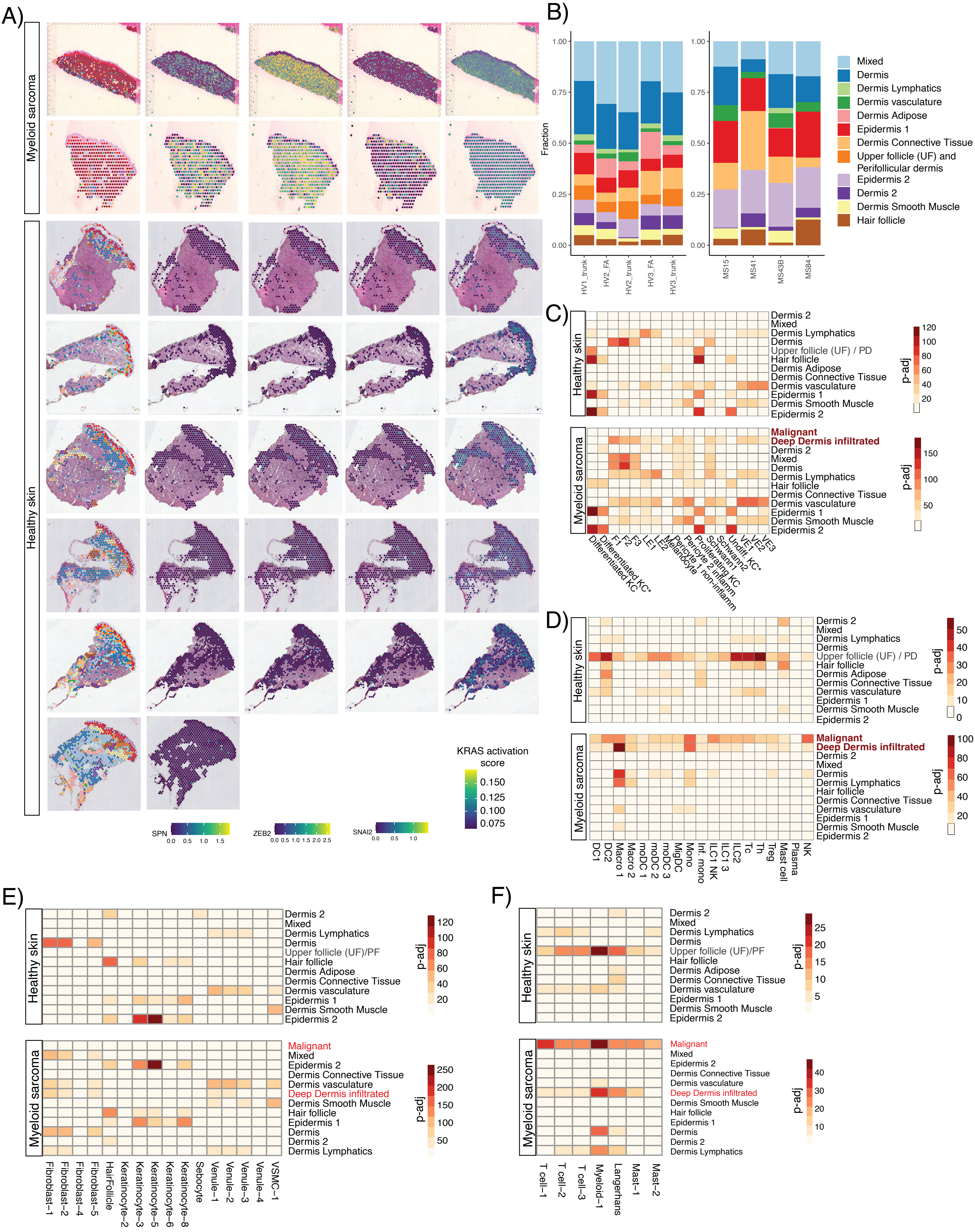

### Supplementary Figure 8

Extended Data Figure 8. RAS inhibition in a murine model of myeloid sarcoma

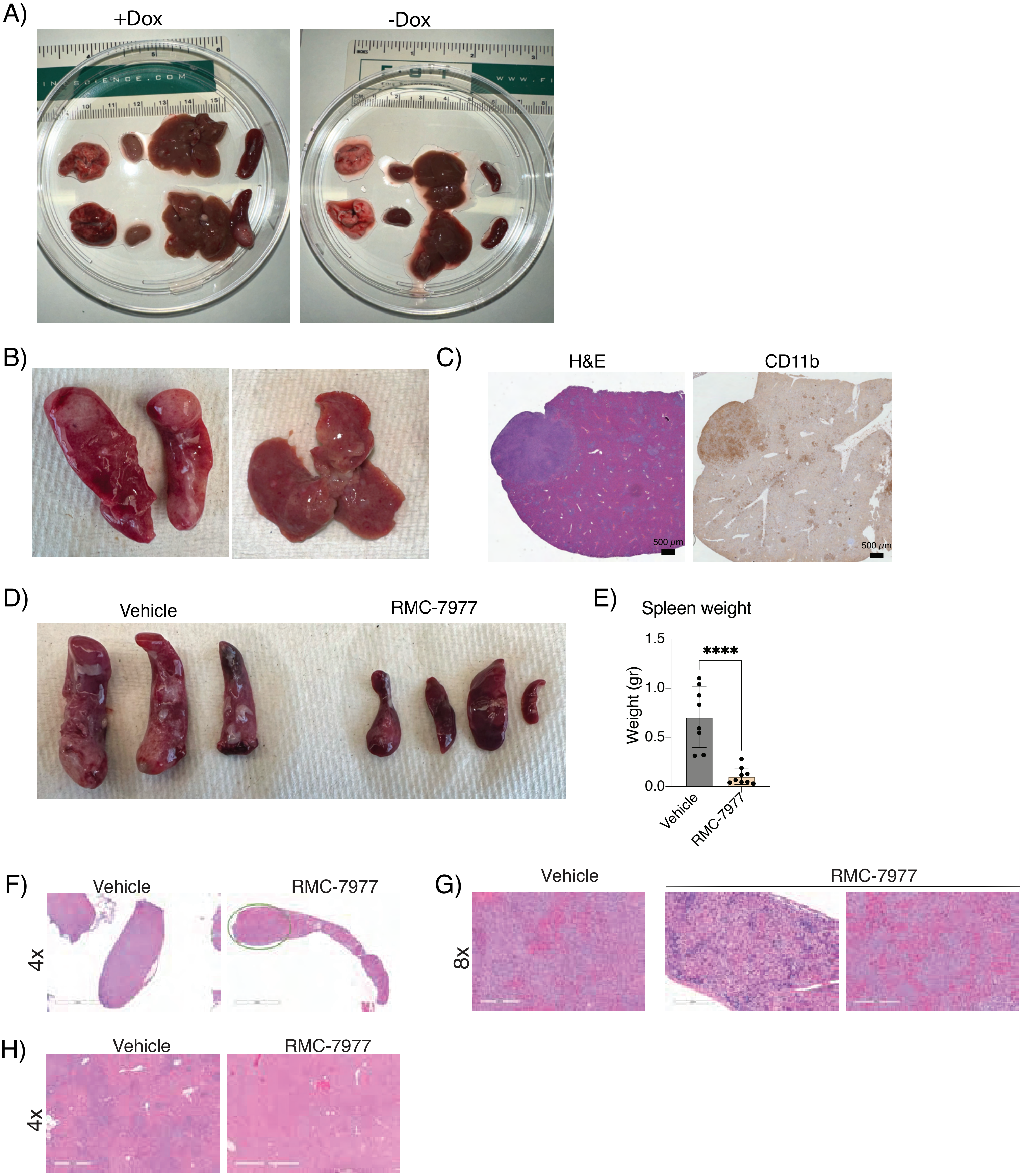
