## Supplementary Figure 2 for "Multiomic characterization, early detection, and therapeutic targeting of myeloid sarcoma"

Extended Data Figure 2: Myeloid sarcoma clones exist at low VAF in the BM and show site-specific evolution.

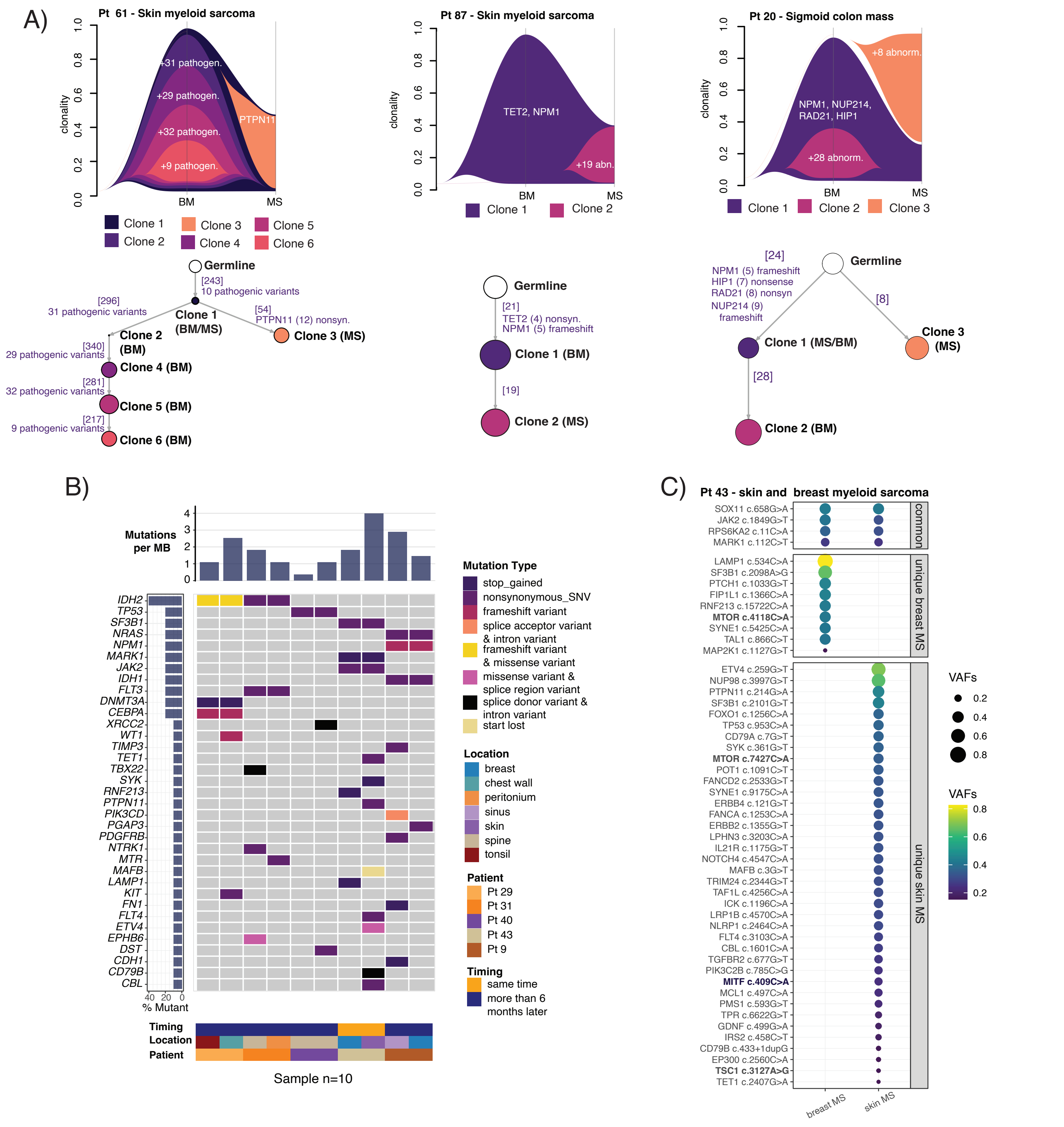
