## Supplementary Figure 3 for "Multiomic characterization, early detection, and therapeutic targeting of myeloid sarcoma"

Extended Data Figure 3: Circulating peripheral blood blasts resemble bone marrow disease in myeloid sarcoma patients.

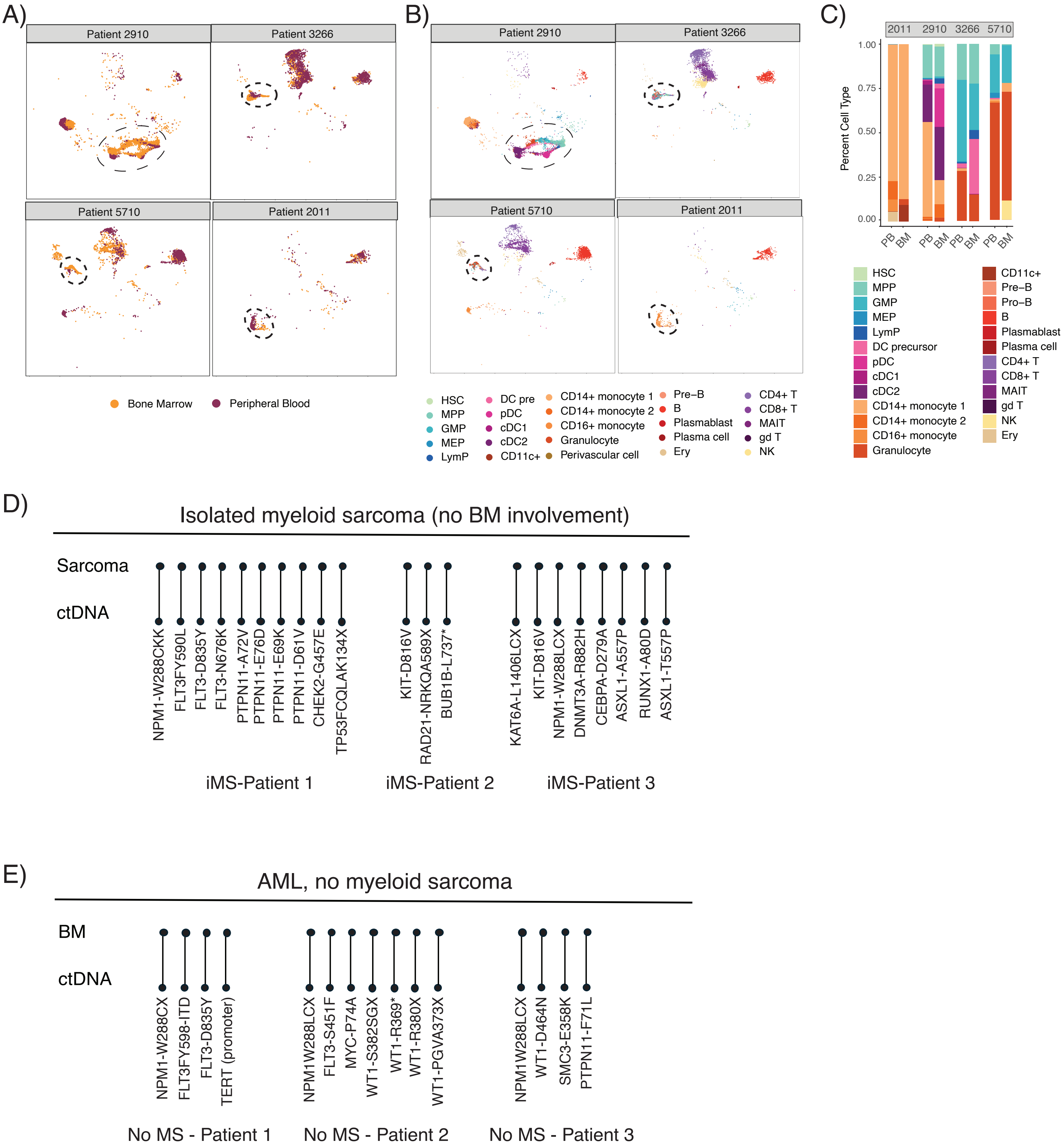
